# Engineered collagen-coated scaffolds for tendon regeneration: a multifunctional drug delivery approach

**DOI:** 10.1101/2025.05.25.655996

**Authors:** Chiara Di Marco, Francesca Romano, Francesco Lopresti, Simona Campora, Albert Comelli, Roberto Di Gesù, Vincenzo La Carrubba

## Abstract

Tendon injuries rank among the most common musculoskeletal disorders, with their incidence rising steadily due to increased sports participation and an aging population. Current surgical treatments often fall short of clinical expectations due to poor biocompatibility, suboptimal mechanical properties, and frequent post-surgical fibrosis. Tissue engineering, however, offers promising alternatives by using biocompatible scaffolds that mimic the native tendon structure. This study aimed to develop an advanced tendon-like bundle replicating the microarchitecture and mechanical strength of native tendons. The scaffold was constructed from aligned electrospun poly-lactic acid (PLA) microfibers and designed to function as a drug delivery system for Rolipram, an antifibrotic agent. To improve cell-scaffold interactions, the scaffolds were coated with type-I collagen, a primary component of the tendon extracellular matrix (ECM). Morphological analyses confirmed the successful fabrication of smooth, well-aligned, bead-free fibers with controlled diameters, closely resembling the natural orientation and dimensions of tendon collagen fibers. Drug release was monitored for both uncoated and collagen-coated electrospun mats, with no burst release observed. Histological analysis demonstrated effective cellular infiltration by human tenocytes, with cells distributed throughout the scaffold after 14 days in culture. These results underscore the potential of tendon-like scaffolds as a promising platform for tendon repair, setting the stage for further optimization.

## INTRODUCTION

More than 1.7 billion musculoskeletal diseases are reportedly hitting population yearly worldwide, and nearly 50% of those diseases are classified as tendon injuries (TI) [1].

TI encompasses a wide range of pathological conditions, including ruptures, overuse injuries, and inflammatory or degenerative disorders such as tendinopathies [2].

Moreover, extensive, or end-stage TI may have a significant clinical impact as they negatively influence the patients’ quality of life, limiting their ability to conduct daily activities, and often affecting also their social life. [2]

The key role that tendons play within the musculoskeletal system is directly linked to such a significant clinical impact of TI. Indeed, tendons are anatomic components capable of transmitting forces from muscles to bones, ultimately allowing the movement around a joint *in vivo* [3]. This is possible thanks to their high-ordered hierarchical structure based on collagen molecules arranged in fibrils grouped in fibril bundles, fascicles, and fiber bundles that are parallel to the longitudinal axis of the tendon [4]. Unfortunately, such a complicated architecture negatively affects the tendons resistance, which decreases over time following repeated stress and overloading, ultimately leading to microtrauma that very often causes tendon ruptures (TRs). This is particularly evident for high-bearing tendons such as the Achilles tendon, which is among the most frequently injured tendons in the body [3].

Regardless of the tendon affected, TRs represent a relevant unmet clinical need due to the limited self-healing capacity of tendons, mainly due to their low vascularity and cellularity [5]. Currently, the surgical repair is the gold standard for the TRs treatment able to re-establish the original tendon’s structural integrity. Sadly, such structural integrity often does not correspond to a tendon’s functional restoration as the surgical repair may induce the formation of nonfunctional fibrotic tissue, which is unable to support mechanical stresses in the joints [6], [7].

In this complex scenario, tissue engineering (TE) approaches have gained importance in clinical practice, both to overcome surgery complications and to develop approaches for early-stage TIs. Electrospinning (ES) is a widely adopted technique for producing 3D scaffolds for TE applications. It allows precise control over nano/micro architectures, enabling the fabrication of aligned meshes, bundles, and layered or braided structures." [8]. These kinds of architectures typically show conspicuous porosity, and a high air-to-matrix ratio, which maximize the surface available within the scaffolds. For this reason, electrospun scaffolds have been extensively employed also as drug delivery systems (DDS) with high tunable kinetics release, which may be controlled by modifying the scaffolds’ microarchitecture (e.g.: fibers diameters, matrix hydrophobicity) [9],[10] [11], [12].

This feature may be exploited to address the drawbacks of surgical approaches to TIs, such as bacterial infections [13], pathological inflammatory responses [9] or fibrosis [9], [14]. Remarkably, tendon fibrosis (TF) also occurs after spontaneous healing of tendon tears [15], leading to partially nonfunctional tendinous structures. TF is mainly due to a Phosphodiesterase (PDE) misregulation, which is reportedly linked to fibrotic responses *in vivo* [16], ultimately triggering inflammation and hypertrophy. Specifically, two of the PDE4 subfamily isoforms PDE4A/B are directly involved in TF, and tendons hypertrophy [17]. Today, several pharmacological approaches to fight fibrosis have been explored including the use of Rolipram, a powerful PDE4A/B inhibitor studied since the late 90’s [17]. Several studies have demonstrated that inhibiting PDE activity can produce antifibrotic effects. Specifically, increased PDE4A and PDE4B expression in hypertrophic ligamentum flavum suggests a role in its hypertrophy. Rolipram, a PDE4 inhibitor, has shown strong antifibrotic effects, primarily by suppressing TGF-β1 activity through the restoration of normal (extracellular signal-regulated kinase) ERK1/2 signaling pathways [18].

With this paper we aim to describe our multimodal approach based on the combination of TE, and drug delivery, to treat TFs triggered by surgical repair of tendon tears. In our work, we fabricated an ES structure with a tendon-like microarchitecture, capable of acting as a drug delivery system for Rolipram. The main idea is to fabricate a scaffold able to support both structurally, and functionally, the tendon regeneration after the surgical repair of tendon tears.

The scaffolds are made of aligned poly-lactic acid (PLA) nanofibers coated with type I collagen to enhance the overall biological compatibility, as well as to slow down the Rolipram early-phase release. We used a 3D-printed bundle-maker developed in house to reproduce a tendon-like architecture at the mesoscale in a reproducible manner. The drug release profile was measured An accurate analysis of the Rolipram release profile allowed us to quantify the drug delivered over time. Moreover, we used human primary tenocytes (hTCs) to cellularize the scaffold for a preliminary cell adhesion investigation. Interestingly, we observed a slower drug release from scaffolds collagen-coated and an improvement in the mechanical properties in dry and wet environments.

## MATERIALS AND METHODS

PLA (200D) was supplied by NatureWorks (Minneapolis, USA) while chloroform (TCM), acetone (Ac) (ACS reagent, > 99.5 percent), dimethyl sulfoxide (DMSO), Triton X-100, and Rolipram were from Sigma-Aldrich (St. Louis, USA). Cell culture supplies were purchased from Euroclone (Pero, Italy): fetal bovine serum (FBS) South America, Penicillin/Streptomycin solution (P/S), 1x Trypsin/ethylenediaminetetraacetic acid (EDTA) in PBS, and GlutaMAX®. Dulbecco’s Modified Eagle Medium (DMEM) was from WVR (Radnor, USA). Sodium pyruvate (NaPyr) and T175 tissue culture flasks were purchased from Corning (Corning, USA). Human PDGF-_BB_ and Recombinant human TGF-β_3_ were from Peprotech (Cranbury, USA). Prolene polypropylene threads from Ethicon (Somerville, USA) were used for suturing the scaffolds. SW-1353 cells were purchased from American Culture Type Collection (Manassas, USA), while cryopreserved primary hTCs from ZenBio (Research Triangle Park, USA). Healthy TCs from three different donors were used in the experiments; their demographic data are shown in Table 1. For immunostaining, Mouse and Rabbit Specific HRP/DAB Detection IHC kit (ab64264), mouse anti-human Tenascin-C (ab86182), rabbit anti-human SOX9 (ab185966), rabbit anti-human Collagen I (ab233080), rabbit anti-human Tenomodulin (ab203676), Donkey anti-Mouse DyLight 550 (ab49945), Donkey anti-Rabbit DyLight 488 (ab96919), and Donkey anti-rabbit DyLight 594 (ab96921) antibodies were purchased from Abcam (Cambridge, UK); tween-20 and Bovine serum albumin (BSA) were purchased from Fisher Bioreagents (Pittsburg, USA).

**Table 1.** hTCs donor demographic data.

| Donor number | Lot | Age (years) | Gender | Ethnicity | Source |
| --- | --- | --- | --- | --- | --- |
| 1 | TEN042921A | 59 | M | Caucasian | Achilles tendon |
| 2 | TEN012121A | 87 | M | Caucasian | Achilles tendon |
| 3 | TEN030817E | 96 | M | Caucasian | Achilles tendon |

### Preparation of tendon-like scaffolds

#### Electrospinning

The electrospinning process was conducted using a semi-industrial electrospinner (NF-103, MECC CO., Ltd., Japan) configured vertically. A 10% w/w polymeric solution of PLA was prepared in a 2:1 v/v mixture of TCM and Ac. The process parameters were optimized as follows: a flow rate of 0.8 mL/h, a needle-to-collector distance of 17 cm, an applied voltage of 17 kV, and a stainless-steel needle (19-G) was used as spinneret. To produce aligned fibers, a grounded rotary drum collector with a length of 20 cm and a diameter of 10 cm was used and set with a speed equal to 3000 rpm. The electrospinning procedure was carried out for 180 minutes, resulting in a PLA electrospun membrane with an approximate thickness of 40 μm.

In the fabrication of drug-loaded scaffolds, Rolipram was incorporated at a concentration of 500 mmol/L (500 mM), dissolved in DMSO following the procedure reported by Tokuara et al. [19] , to achieve a drug-to-polymer weight ratio of 4.6 µg/mg. To determine the appropriate volume of drug solution to be added, 1.885 g of PLA were dissolved in 15 mL of chloroform/acetone mixture (2:1 v/v), to obtain a 10% w/w polymer solution. Based on the desired drug loading, 8.671 mg of Rolipram were required, corresponding to 63 µL of the 500 mM solution. DMSO was selected for drug solubilization due to its due to its high solvating capacity and compatibility with the organic solvents used during the electrospinning process. All electrospinning parameters were maintained identical to those employed for electrospun mats without drug incorporation. The resulting drug-loaded scaffolds were stored at 4 °C until needed further use and are referred to as “PLA/Rol” throughout the study.

#### Collagen Coating

The PLA/Collagen electrospun membranes were obtained by placing a coating of collagen on the fibers’ surface.

Type I collagen was extracted from rat tail tendons by optimizing the protocol described by Rajan and colleagues [20]. Initially, rat tails were washed in sterile water containing 500 units/mL penicillin G and 500 μg/mL streptomycin (Euroclone, Celbar). Afterward, the tails were dried and surgically incised to remove the skin, isolating the white collagen fibers from the tendons. These fibers were collected and placed in phosphate-buffered saline (PBS) containing 500 units/mL penicillin G and 500 μg/mL streptomycin (Euroclone, Celbar). Next, the collagen fibers were sequentially washed in acetone for 5 minutes, 70% isopropanol for 5 minutes, and 0.02 N acetic acid under agitation for 48 hours at 4°C. The resulting viscous collagen solution was filtered over flake ice and sterilized by dialysis in cold 1% chloroform. Subsequently, it was dialyzed in 0.02 N acetic acid, and its concentration was determined by spectrophotometric analysis using a microplate reader (Synergy HT, Biotek, Winooski, VT, USA) at a wavelength of 280 nm, calibrated against a commercial collagen standard curve. A solution of 4 ml of Collagen I/water/AA was placed in 0.1 g of electrospun PLA rectangle-shaped until it was completely absorbed. Then, scaffolds were left dry under a fume hood overnight at room temperature. PLA mats that underwent surface functionalization with collagen are labeled as “PLA/Coll” and “PLA/Rol/Coll”.

The weight fraction of collagen within the scaffold was determined by subtracting the scaffold weight prior to coating from its weight after coating and normalizing this difference to the final (post-coating) scaffold weight. Assuming that the entire weight increase is attributable to collagen, an average collagen weight fraction of 28% was determined for both PLA/Coll and PLA/Coll/Rol systems.

#### Fabrication of tendon-like scaffolds

A tendon-like structure was obtained from a 1 x 2.5 cm rectangle-shaped scaffold. The scaffold was then modeled using a 3D-printed custom-made bundle-maker in order to obtain a bundle-like architecture that mimics the in vivo structure of tendons. The fabrication process involved several steps: first, the rectangular scaffold was secured between the clamps of the bundle maker. Next, a crank mechanism was rotated, causing the clamps—and consequently the scaffold mat itself—to twist into a bundle formation. At the end of the process, the bundle ends were secured with a polypropylene suture thread, and tied into a loop knot.

### Scaffold characterization

#### Scanning electron microscopy (SEM) analysis

The fibrous mats’ morphology was examined using a scanning electron microscope (SEM) (FEI QUANTA 200, FEI Company, Hillsboro, USA). Before imaging, the samples were gold-coated for 60 seconds using a sputter coater (Scancoat Six, Edwards Laboratories, Milpitas, USA) in an argon atmosphere to minimize electrostatic discharge during imaging. The resulting images were analyzed with ImageJ software to determine the fiber diameter distribution. For each sample, the measurements were averaged over 100 data points, derived from three distinct images.

#### Fourier-transformed infrared with attenuated total reflectance (FTIR-ATR) spectroscopy

Fourier-transform infrared spectroscopy with attenuated total reflectance (FTIR-ATR) was employed to analyze the chemical surface properties of the samples and verify the presence of collagen on the PLA electrospun scaffolds. This technique allowed us to identify specific functional groups and confirm the successful surface modification of the scaffolds. Each sample underwent spectral scanning across the wavenumber range of 4000 to 500 cm[¹, with a resolution of 4 cm[¹. The analysis was carried out using an IRTracer-100 spectrophotometer (Shimadzu, Milan, Italy), which provided high-resolution spectra suitable for assessing both the polymeric matrix and the collagen coating, with 2 samples analyzed for each condition.

#### μ-CT analyses

Micro-computed tomography (µCT) scans were performed to analyze the 3D structure of the fibrous bundle-shaped scaffolds. CT datasets were acquired using a preclinical µCT (SkyScan microtomography 1275) (Bruker Company - Brussels, Belgium). The scaffold was imaged using 1709 views in a high-resolution configuration, a field-of-view (FOV) of 4096 × 4096, X-ray energy of 35 kV, a current of 100 μA, and the size of each CT-voxel was equal to 21.952 µm^3^ (2.8 × 2.8 × 2.8 μm). The DICOM (Digital Image Communications in Medicine) images were obtained using a modified Feldkamp algorithm. The 3D rendering of the sample was performed using a 3DSlicer V. 5.6.2 (Earth, USA).

#### Mechanical testing

The mechanical properties of the electrospun scaffolds were assessed using a universal testing machine (UTM, model 3367, Instron, Norwood, MA, USA), equipped with a 1 kN load cell and a BioPulse bath for environmental control. Rectangular samples measuring 10 × 70 mm, with a thickness of approximately 40 μm, were prepared by cutting the membranes along the fiber alignment direction (radial to the collector). The mechanical tests were performed under two distinct conditions: in a "dry" state at room temperature and in a "wet" state with samples submerged in a water bath maintained at 37°C to simulate physiological conditions. The tests were conducted with a crosshead speed of 5 mm/min and a gauge length of 30 mm. For each sample, stress-strain curves were generated, and representative curves were analyzed to evaluate mechanical performance. The elastic modulus was calculated from the linear region of the stress-strain curves, reflecting the stiffness of the materials.

A total of three specimens were tested for each condition, and the mechanical parameters, including elastic modulus, were averaged and reported along with their standard deviations to ensure reliability and statistical accuracy.

#### UV-Visible spectroscopy

The release profile of Rolipram from PLA/Rol and PLA/Rol/Coll scaffolds was investigated over time in phosphate-buffered saline (PBS) at 37 °C using a Cary 100 Scan UV-Vis spectrometer by Varian (Palo Alto, USA). The characteristic absorption peak of Rolipram was monitored at 280 nm. The experiment was conducted under these conditions to simulate physiological temperature. To enable quantification a calibration curve was constructed by measuring the absorbance of Rol/PBS solutions across a concentration range of 0.0015-0.6 mg/mL (Fig. S1). Rectangular samples of PLA/Rol and PLA/Rol/Coll (approximately 4 mg each) were immersed in 1.5 mL of pre-warmed PBS and incubated at 37 °C. At each sampling interval, the absorbance of the release medium was measured, and the samples were transferred to fresh containers with 1.5 mL of new PBS solution at 37 °C to continue the release process. The cumulative drug release was calculated by summing the amount of Rolipram released at each time point. Three replicates were analyzed per sample type, and the average values were reported.

Data were also reported as M_t_/M∞ Vs time where, M_t_ is the cumulative amount of Rolipram released at time t, while M∞ represents the total amount of drug initially loaded into the scaffold, assumed to be the maximum releasable quantity. To elucidate the diffusion behavior and release kinetics of Rolipram, the experimental data were fitted using a power-law equation (Equation 1):

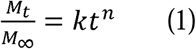

Where t represents the release time, k is a kinetic constant associated with the rate of release, and n is the diffusion exponent which provides insight into the release mechanism, according to Peppas [21]. Since the Peppas model is typically applicable only to the initial release phase, where M_t_/M∞ is lower than 60% [22], the fitting for parameter extraction was performed by analyzing the experimental data within the first four hours for both PLA/Rol and the PLA/Rol/Coll systems. Within this time window, the cumulative release remained below 60%, thereby satisfying the model assumptions and allowing for a reliable estimation of the *k* and *n* values that accurately describe the early-stage release kinetics.

### Cell culturing

#### 2D tenocytes culturing

hTCs (ZenBio) at passage 3 were seeded in T175 tissue culture flasks (10^6^ cells/cm^2^) and maintained in 10% v/v Fetal Bovine Serum (FBS) + 1% v/v Penicillin/streptomycin (P/S), 1% v/v GlutaMAX®, 1% v/v Sodium pyruvate (NaPyr), until reaching 80% confluence. Cells were passaged using 0.1% Trypsin/ EDTA and grown in a humified incubator at 37° C and 5% CO_2_ by Thermo Fisher Scientific (Waltham, USA), replacing the growth medium every second day. To study the impact of different growth media on proliferation kinetics and phenotype maintenance, passage 5 hTCs were seeded at 10^4^ cells/cm^2^ in 24-well plates for 14 days in five different growth media, as detailed in Table 2. Each condition was tested in triplicate.

**Table 2.**
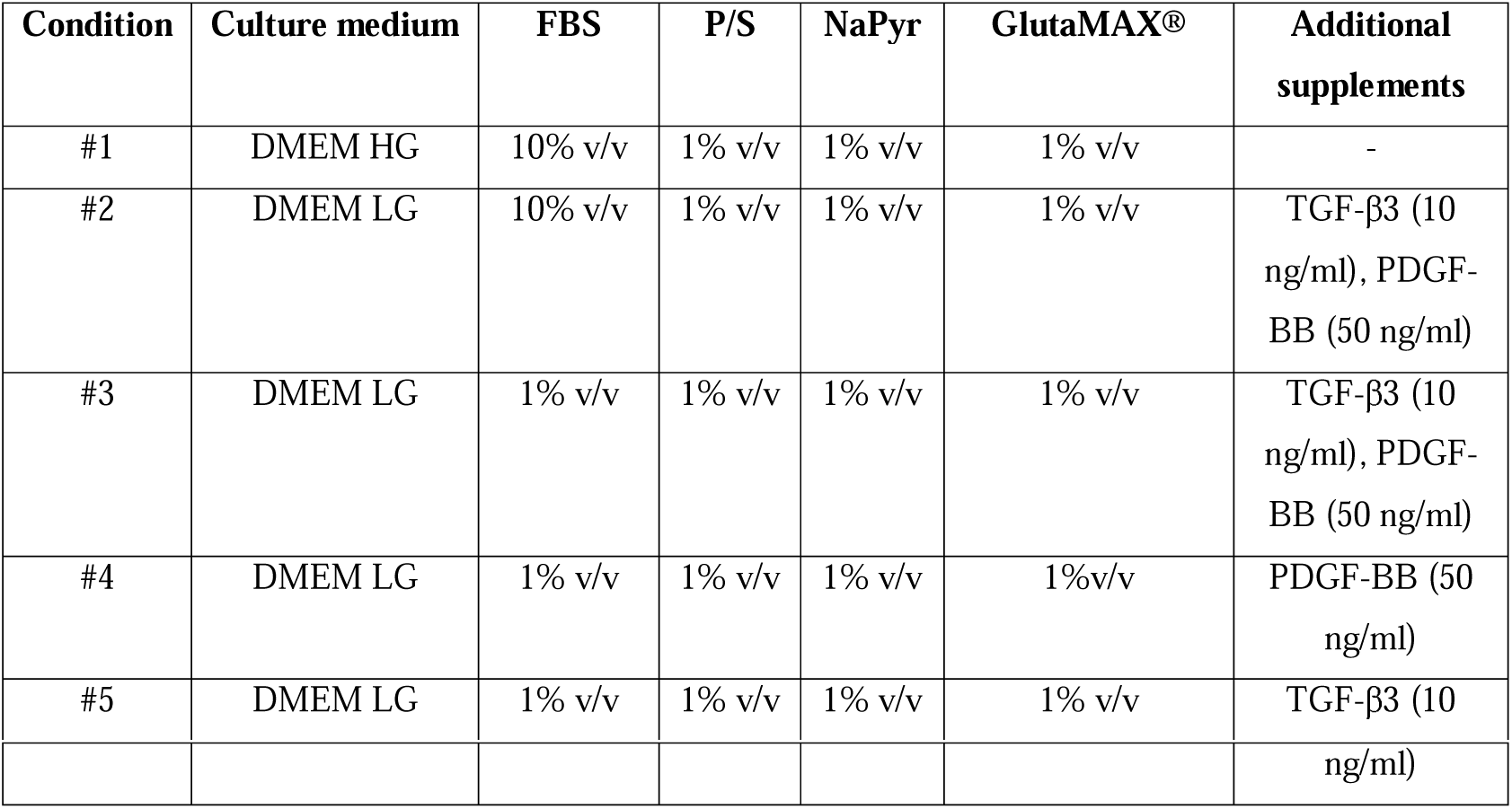
Summary of growth media formulations used for 2D hTCs culture.

Immunocytochemistry was performed to assess the expression of the tenogenic marker Tenascin-C (TNC) and SOX9 (a transcription factor that regulates chondrogenesis), after 14 days of culture. The selection of TNC as a positive marker and SOX9 as a negative marker was intended to evaluate a potential transdifferentiation towards the chondrogenic lineage. Following cell culture, wells were washed with PBS and fixed with 4% w/v paraformaldehyde (Sigma-Aldrich) at RT for 30 min. After washing the samples three times with PBS, cells were permeabilized with 0.2 % v/v Triton X-100 for 10 min and then washed again with PBS. Cells were subsequently blocked with a blocking buffer, consisting of 3% BSA in PBST (0.05% v/v Tween-20 in PBS) at RT for 1 h. Next, the samples were incubated with a primary anti-Tenascin C antibody (ab86182, 1:50 dilution in blocking buffer) and a primary anti-SOX9 antibody (ab185966, 1:200 in blocking buffer) in each case at 4 °C overnight. After the incubation time, samples were washed three times with PBS and then incubated with the secondary antibodies Donkey Anti-Rabbit DyLight 488 (ab96919, 1:50 dilution) and Donkey Anti Mouse DyLight 550 (ab49945, 1:50 dilution) diluted in the same blocking buffer. The incubation was carried out for 1 hour in the dark. Subsequently, samples were washed 3 times with PBS and stored at 4° C in the dark until visualization. Fluorescence imaging was performed using EVOS M5000 by Thermo Fisher Scientific, a fully integrated, digital inverted benchtop microscope designed for fluorescence microscopy. For each condition, six independent images were acquired. Each image was first converted to 8-bit format using ImageJ software. A uniform threshold was applied across all images to ensure consistency in signal detection. Following thresholding, the mean fluorescence intensity for each image was recorded and employed for subsequent quantitative analysis.

#### 3D tenocytes culturing

For the preliminary cell-adhesion assessment, PLA bundle-shaped scaffolds were cellularized with human chondrosarcoma SW-1353 cells by adding 50 µL of a cell suspension containing 10^5^ cells onto the scaffold. On Day 2 of culture, nuclei were stained with 4’,6-diamidino-2-phenylindole (DAPI) for 20 min at room temperature.

hTCs at passage 5 were used to evaluate cell attachment and viability on the scaffolds. Cells were cultured using DMEM with low levels of glucose supplemented with 10% v/v FBS, 1% v/v P/S, 1% v/v NaPyr, 1% v/v GlutaMAX®, PDGF-_BB_ and TGF-β3. hTCs were then seeded onto the scaffolds by applying 30 µL of cell suspension with 2.5 × 10[ cells to each scaffold (three scaffolds/condition). Scaffolds were incubated at 37°C with 5% CO_2_ and supplemented with the complete growth medium every other day for a total of 14 days. Following a two-week culture period, the samples were fixed in a 4% w/v paraformaldehyde solution for 30 minutes and then washed three times with PBS. Next, samples were embedded in an optimal cutting temperature (OCT) compound and immediately frozen. The orientation during mounting was carefully adjusted to ensure that the scaffold blocks were cut either longitudinally or transversely. These samples were subsequently sectioned into 4 μm thick slices using a cryostat and transferred onto Knittel Glasbearbeitungs GmbH glass slides (Braunschweig, Germany). Cryosectioned scaffolds were subjected to immunofluorescence staining following a standard protocol. Samples had previously been fixed in 4% paraformaldehyde. Antigen retrieval was performed by immersing the slides in 10 mM sodium citrate buffer and heating them in a microwave oven at 90 °C for 15 minutes. After heat-induced epitope retrieval, the slides were allowed to cool at room temperature for 30 minutes, followed by three washes with PBS, each lasting 5 minutes. To permeabilize the samples, slides were incubated in 0.1% (v/v) Triton X-100 prepared in Tris-buffered saline (TBS) for 10 minutes at room temperature. After a brief PBS rinse, slides were air-dried and then delimited using a hydrophobic barrier pen to prevent reagent mixing across samples. Blocking was performed using Mouse and Rabbit Specific HRP/DAB Detection IHC kit (ab64264) for 45 minutes at room temperature. Rabbit anti-human Collagen I (ab233080, 1:50 dilution) and rabbit anti-human Tenomodulin (ab203676, 1:50 dilution), both diluted in the previously described blocking buffer were applied. Slides were incubated overnight at 4 °C in a humidified chamber. The next day, samples were washed three times with PBS (5 minutes each), followed by incubation with the secondary antibody Donkey anti-rabbit DyLight 594 (ab96921, 1:50 dilution), for 1 hour at room temperature in the dark to protect from light-induced photobleaching. After a final 5-minutes PBS wash, slides were gently dried and mounted using a limonene-based mounting medium. Samples were stored in a humidity slide chamber until imaging. For each scaffold, five independent images were acquired for both Collagen I and Tenomodulin (TNMD) with confocal microscope. Image processing was carried out following the same protocol detailed above.

Separate sections obtained from the same cryosectioning procedure were stained with Hematoxylin and eosin (H&E) staining.

### Statistical analysis

Statistical analyses were conducted to evaluate the significance of experimental findings. Drug release data were analyzed using two-way ANOVA, followed by Tukey’s post hoc test to assess pairwise differences between experimental groups. For the 2D immunofluorescence quantification of TNC and SOX9 expression, the non-parametric Kruskal–Wallis test was applied. In the case of 3D immunofluorescence data for Collagen I and TNMD, statistical analyses were carried out using the Mann–Whitney test. In all analyses, a p-value <0.05 was considered as the threshold for statistical differences. Statistical analysis was performed using Prism (GraphPad Software, version 10.4.1).

## RESULTS

### Fibrous alignment in 3D bundle-like scaffolds reproduces native tendon architecture

The 3D tendon-like scaffold fabricated using our custom-made bundle-maker device (figure 1A) demonstrated structural integrity at the mesoscale, with no signs of delamination or structural rupture observed after fabrication and suturing (Figure 1B). Remarkably, the scaffold maintained its structural integrity throughout the entire experimental period in a water-based culture medium (figure 1C).

**Figure 1.**
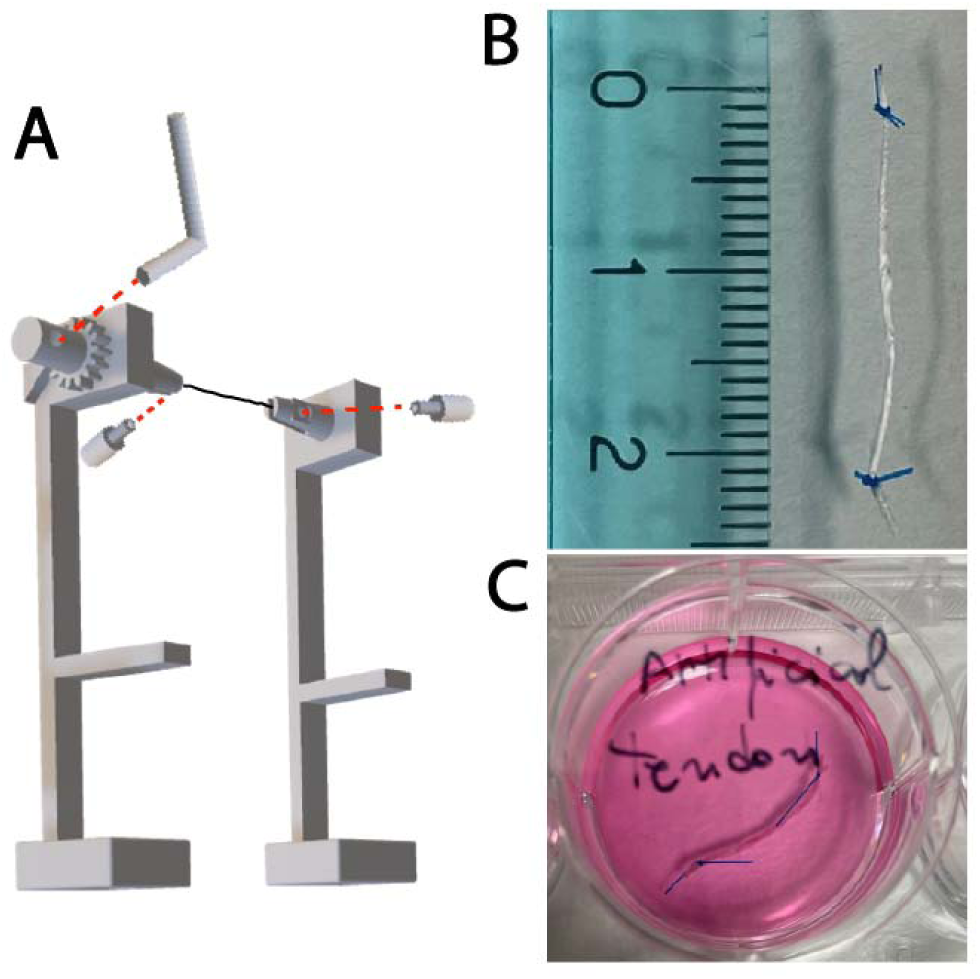
Exploded 3D model of the custom-made bundle-maker used to fabricate tendon-like scaffolds (A). Top view the bundle-shaped scaffold (B). Cellularized scaffold cultured within a 24 well multiwell plate (C).

Such an integrity at the mesoscale corresponds to an anisotropic architecture at the microscale, either for 2D (figure 2), or 3D scaffolds (figure 3), suggesting that our fabrication approach efficiently produced aligned microfibers in each condition. Furthermore, it is worth underlining the characteristic fibers self-assembly in collagen-coated scaffolds (figure 2 B, D), which closely reproduce the hierarchical architecture of native tendons. Conversely, non-collagen-coated scaffolds exhibited a single-fiber microarchitecture characterized by microfibers spatially independent from the nearby surrounding microfibers (figure 2A, C). Such a morphological geometry is coherent with the fibers’ diameter distribution, which increased by 30%, and 15% after the collagen coating of PLA, and PLA/Rol material respectively (figure 2E).

**Figure 2:**
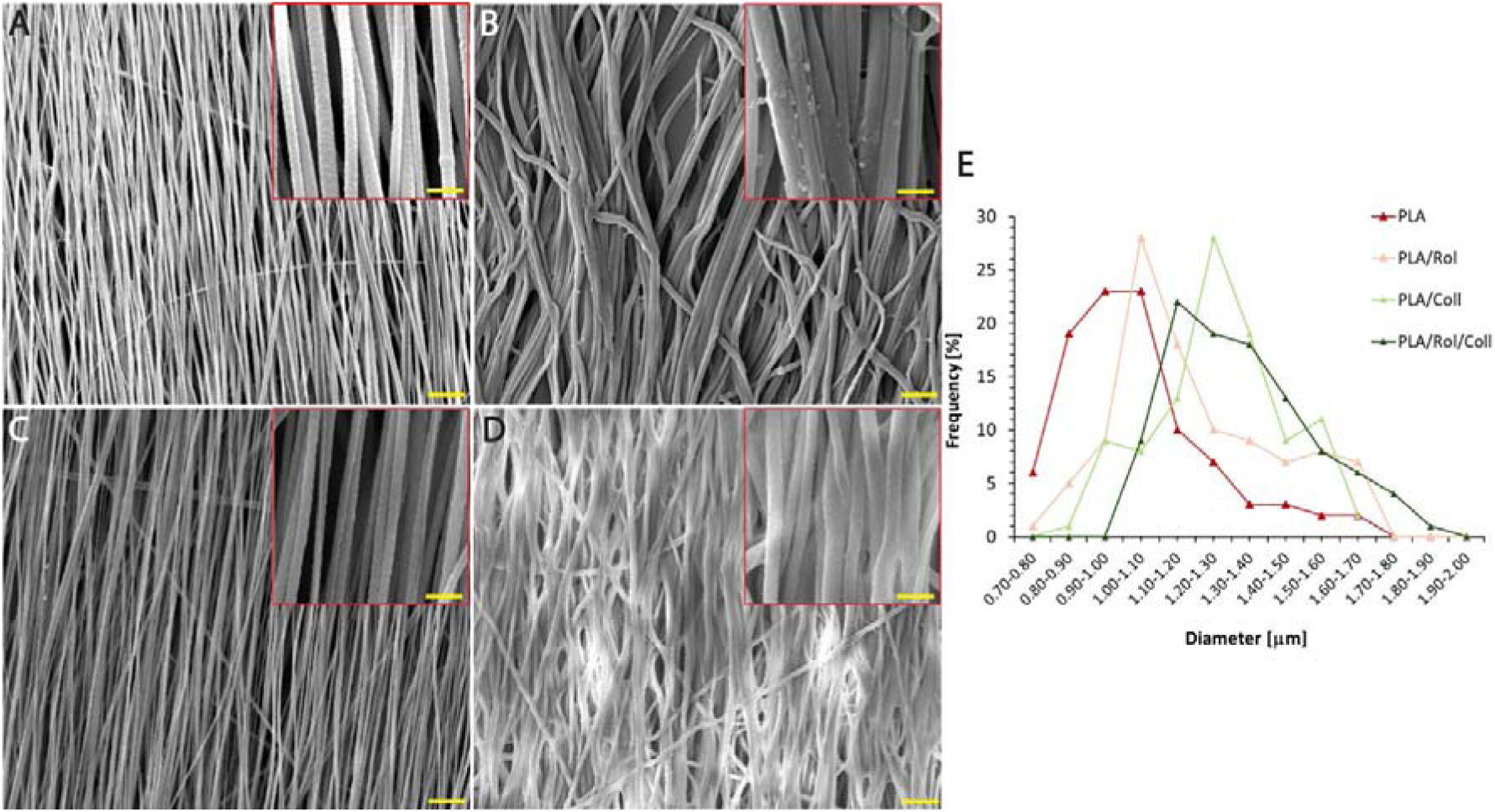
Scanning electron microscopy (SEM) showing fiber morphology of PLA (A), PLA/Coll (B), PLA/Rol (C), and PLA/Rol/Coll (D) electrospun mats. Main scale bar = 10 µm insert scale bar = 2.5 µm. Graphical representation of the fiber diameter distribution (E).

**Figure 3:**
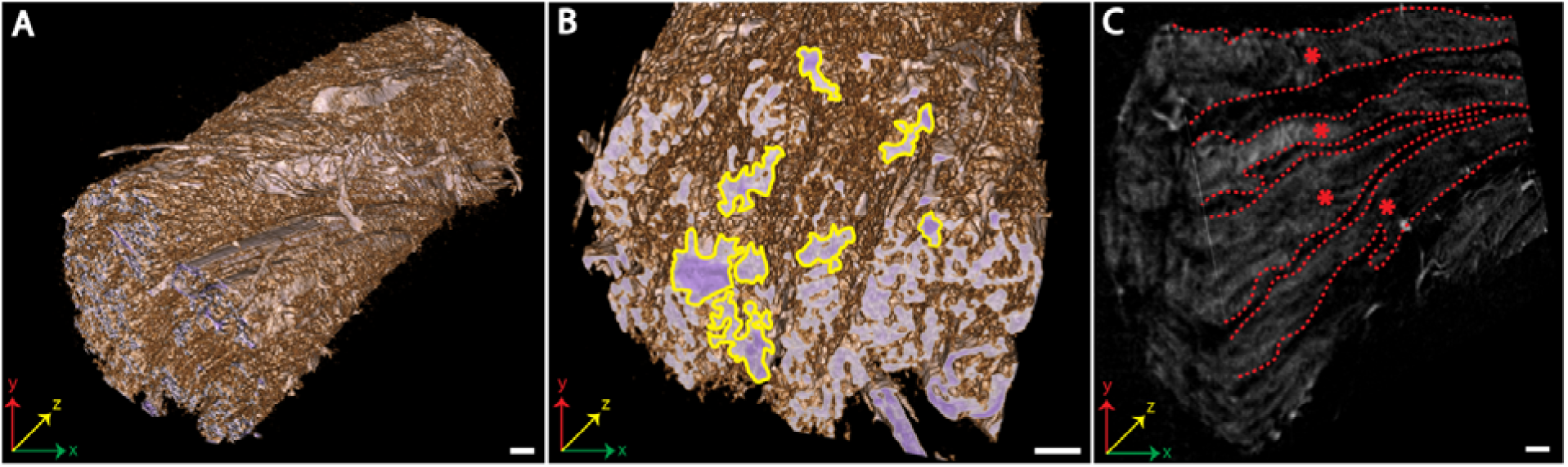
Rendering of µ-CT scans of the bundle-shaped scaffold captured in false colors at different angled views (A, B). X-ray images of a cross-sectional view (C). Yellow areas = fascicular suborganization in transversal view. Red dashed-lines, and red asterisks = extension of fascicular suborganization through longitudinal axis. Scale bar = 100 µm.

Remarkably, the anisotropic microarchitecture was preserved in the tendon-like 3D scaffold as showed by the rendering obtained from µCT multi-scan analyses of a PLA scaffold (figure 3), with a fascicular suborganization geometry similar to that of native human tendinous fibers (yellow areas, figure 3B, red dashed-lines and asterisks, figure 3C) [23].

### Our coating protocol is effective toward a superficial scaffold functionalization

FTIR-ATR analyses performed on 2D ES mats have confirmed the effectiveness of our collagen-coating protocol. Besides strong peaks characteristic of PLA, several absorbing bands coherent with the type I collagen were visible (figure 4). More in detail, narrow, and weak peaks at 2990 cm^-1^, and 2950 cm^-1^ refer to the symmetric, and asymmetric -CH_3_ side moieties stretching respectively [24]. The strong, and narrow peak at 1747 cm^-1^ corresponds to the C=O stretching of the carbonyl group in the main PLA backbone [25]. Moreover, two peaks characteristic of type I Collagen were clearly noticeable at 1650 cm^-1^, and 1550 cm^-1^, corresponding to the C-O stretching of carbonylic Collagen moieties of amide I and II respectively [26] [27]. Any peak related to Rolipram was visible, suggesting a drug distribution within the inner portion of the fibers.

**Figure 4.**
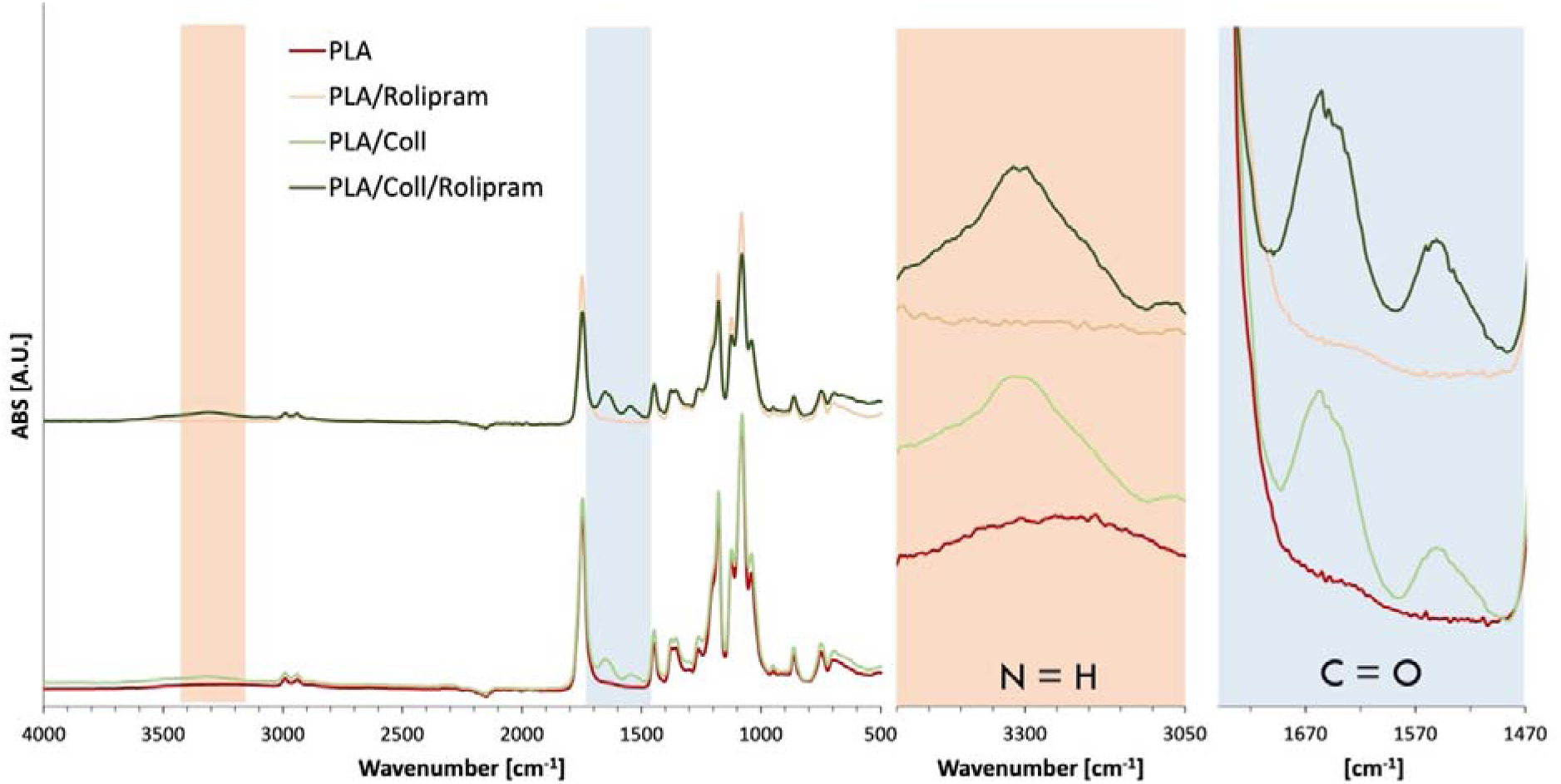
FTIR-ATR spectra of PLA, PLA/Rol, PLA/Coll, and PLA/Rol/Coll electrospun mats.

### Collagen coating enhances scaffold stiffness and supports mechanical integrity in physiological-like conditions

The fibers coated with type I collagen showed to be drastically effective toward the overall stiffness of the scaffolds as described by the behavior of the stress/strain curves either in dry (figure 5A), and in wet conditions (figure 5B). The tensile strength (TS) value was almost doubled in collagen-coated samples in dry conditions, with an increase from 5.36 to 10.54 MPa and from 4.58 to 9.84 MPa for samples with, and without Rolipram respectively (table 3). An analogous behavior was noticeable in wet conditions, with TS values ranging from 3.14 and 8.9 MPa for scaffolds drug-free, and from 2.93 to 9.74 MPa for scaffolds Rolipram-loaded (table 3). The TS variation between coated, and uncoated samples aligns with the trend observed in the elastic modulus (E), which was increased in the coated sample compared to the corresponding uncoated scaffolds both in dry and in wet conditions. More in detail, E increased in coated scaffolds from 102.93 MPa to 140.09 MPa, and from 103.3 to 144.23 MPa for drug-free and drug-loaded scaffolds respectively in dry conditions (figure 6A). A similar trend was visible in wet conditions, where E increased from 54.93 to 132.15 MPa, and from 45.91 to 130.89 MPa for scaffold drug-free, and drug-loaded respectively (figure 6B).

**Figure 5:**
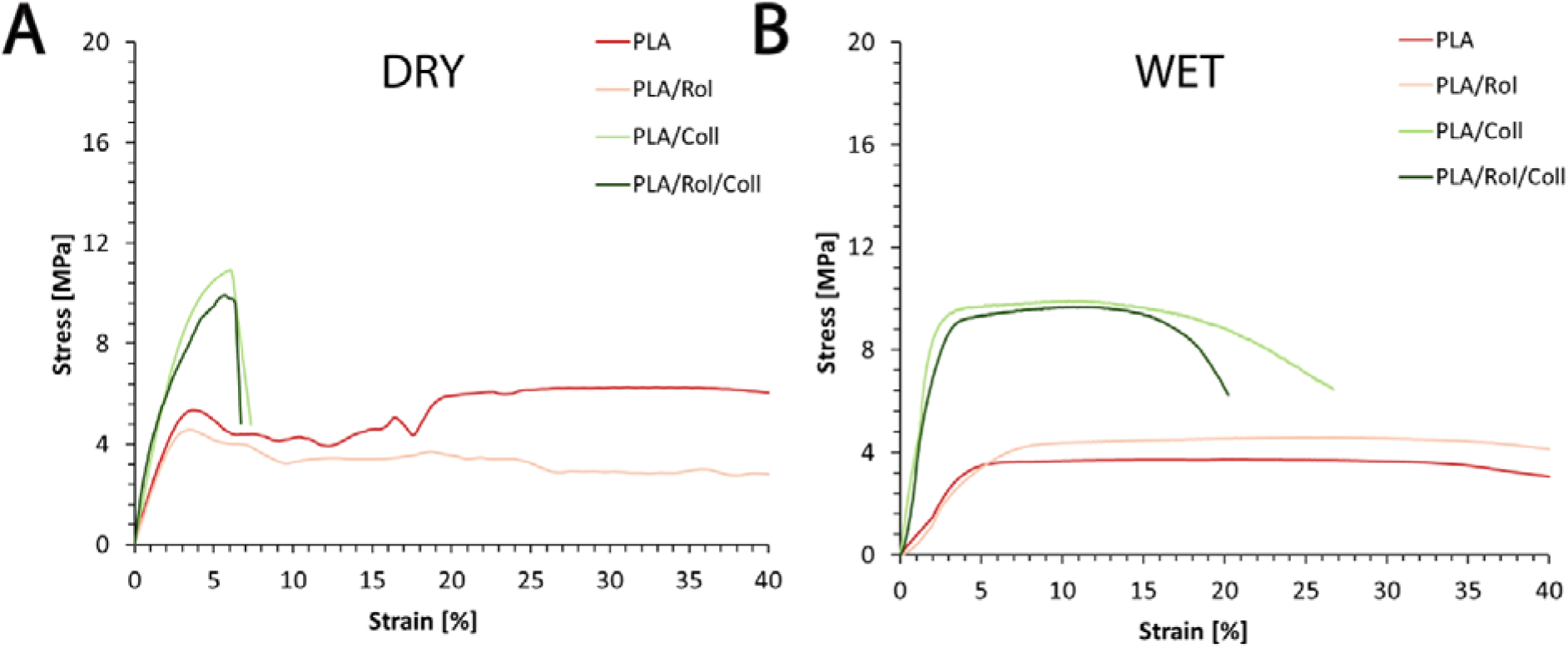
Representative stress/strain curves of PLA, PLA/Rol, PLA/Coll, and PLA/Rol/Coll electrospun mats in dry (A) and wet (B) conditions.

**Figure 6:**
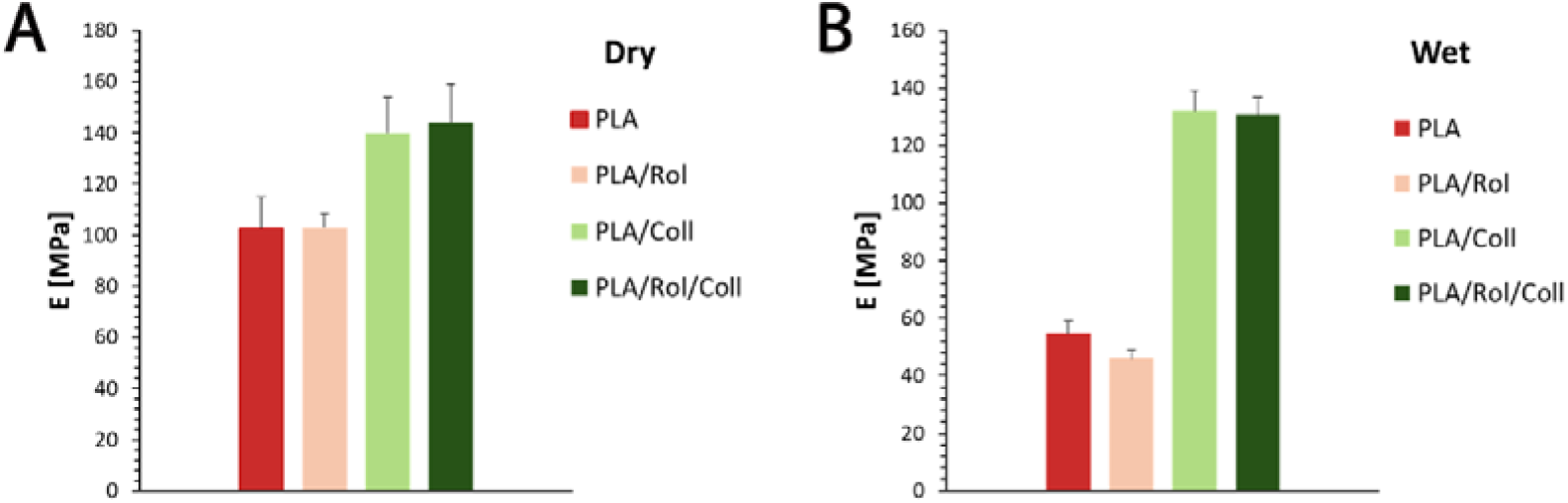
Young’s modulus of PLA, PLA/Rol, PLA/Coll, and PLA/Rol/Coll samples under dry (A) and wet (B) conditions. Data are reported as mean ± SD.

**Table 3:** Tensile properties of PLA, PLA/Rol, PLA/Coll, and PLA/Rol/Coll electrospun mats in dry and wet conditions

| | E [MPa] | TS [MPa] | $\epsilon_t$ [%] |
| --- | --- | --- | --- |
| *Dry condition* |  |  |  |
| PLA | 102.93 ± 12.35 | 5.36 ± 0.42 | 103.82 ± 7.26 |
| PLA/Rol | 103.3 ± 5.16 | 4.58 ± 0.32 | 114.27 ± 13.71 |
| PLA/Coll | 140.09 ± 14 | 10.54 ± 1.26 | 7.32 ± 0.87 |
| PLA/Rol/Coll | 144.23 ± 15.86 | 9.84 ± 0.59 | 6.73 ± 0.47 |
| *Wet condition* |  |  |  |
| PLA | 54.93 ± 4.39 | 3.14 ± 0.34 | 82.46 ± 5.77 |
| PLA/Rol | 45.91 ± 3.67 | 2.93 ± 0.23 | 80.56 ± 7.25 |
| PLA/Coll | 132.15 ± 7.92 | 8.9 ± 0.89 | 27.58 ± 3.03 |
| PLA/Rol/Coll | 130.89 ± 6.54 | 9.74 ± 0.48 | 31.32 ± 1.87 |

Simultaneously, a significant reduction of the elongation at break in coated samples was evident, with a decrease from 103.8% to 7.3% for drug-free scaffolds, and from 114.3% to 6.7% for drug-loaded scaffolds (table 3).

### Collagen-coated scaffolds facilitate a prolonged drug release

The Rolipram kinetics release was thoroughly monitored up to 144 hours from both uncoated (PLA/Rol), and coated (PLA/Rol/Coll) mats (figure 7), referring to a calibration curve built for Rolipram in PBS (figure SI 1). As shown in both panels, PLA/Rol scaffolds exhibited a markedly faster release compared to the PLA/Rol/Coll counterparts. Within the first hour, a significant greater amount of Rolipram was released from uncoated scaffolds (*p* < 0.01), and this trend persisted throughout the entire duration of the experiment. At 24 hours, PLA/Rol had released 3.5 mg of drug per gram of scaffold, whereas PLA/Rol/Coll remained below 2 mg/g. This discrepancy was also evident when data were normalized to the total drug load (M_t_/M∞): uncoated scaffolds released approximately 60% of the loaded Rolipram within the first 4 hours, while the collagen-coated samples required nearly 48 hours to reach the same release fraction.

**Figure 7.**
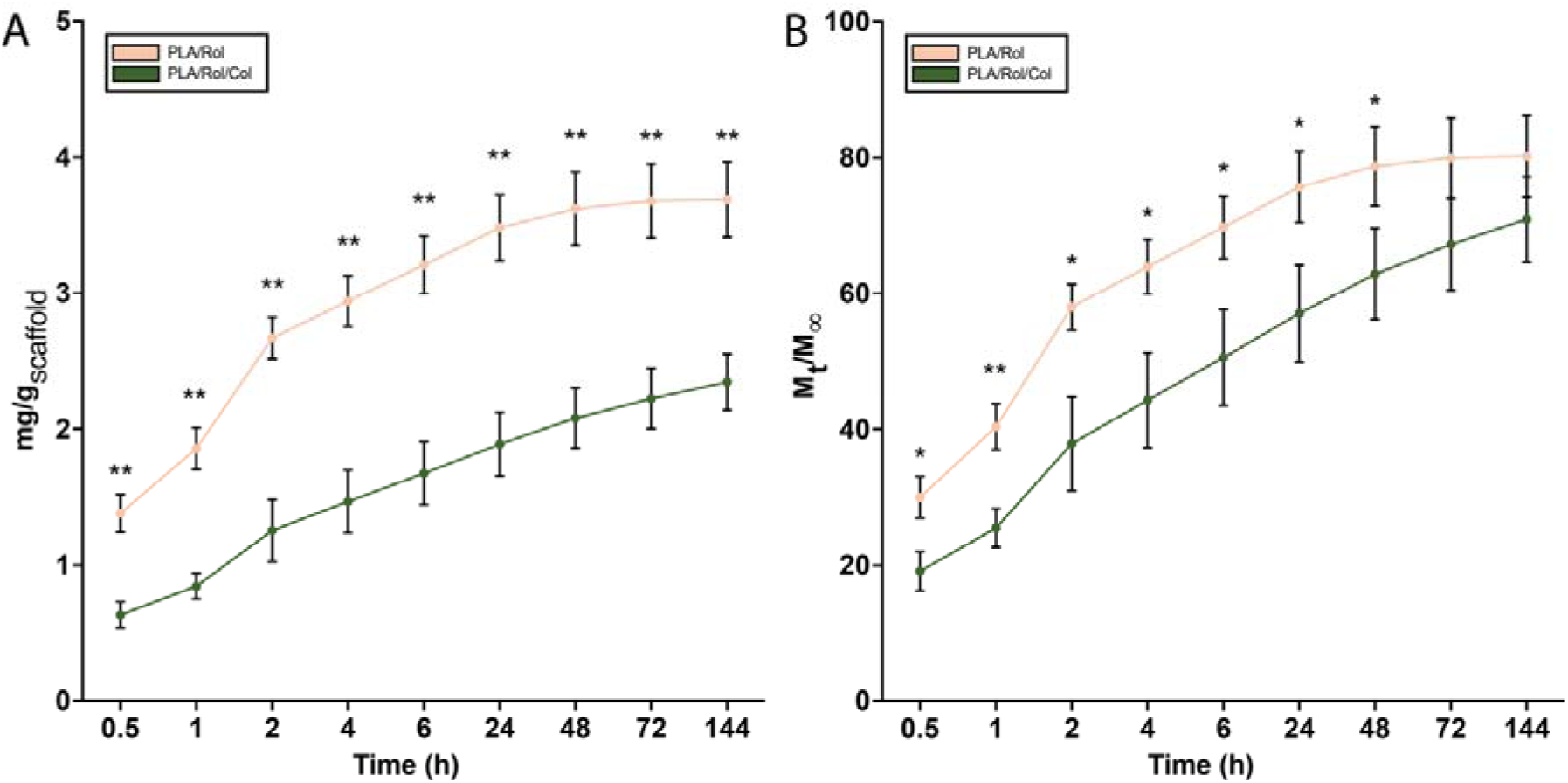
Cumulative release profiles of Rolipram in PBS at 37 °C expressed as A) mg of Rolipram released from 1 gram of scaffold and B) *M_t_/M*∞. Two-way ANOVA followed by Tukey’s post hoc test, * = p < 0.05(*), p < 0.01 (**). Data are reported as mean ± SD (n = 3).

### Optimized medium formulation enhances Tenascin-C expression from hTCs

hTCs easily go toward phenotype modification during monolayer culturing in 2D [28]. Our optimized culturing medium showed to prevent this unwanted phenomenon, preserving the native phenotype of hTCs cultured up to 14 days in 24-well plates using five different media formulations. TCs cultured with medium #2 showed higher secretive activity for Tenascin-C (TNC), which was detected and quantified after immunohistochemical staining (figure 8 I,L). Interestingly, the TNC deposition from TCs cultured within other experimental conditions (#1, #3, #4, and #5) was significantly lower compared to condition #2 (figure 8M). Conversely, we did not observe any significant difference in the production of SOX9 between 5 experimental conditions (figure 8N).

**Figure 8.**
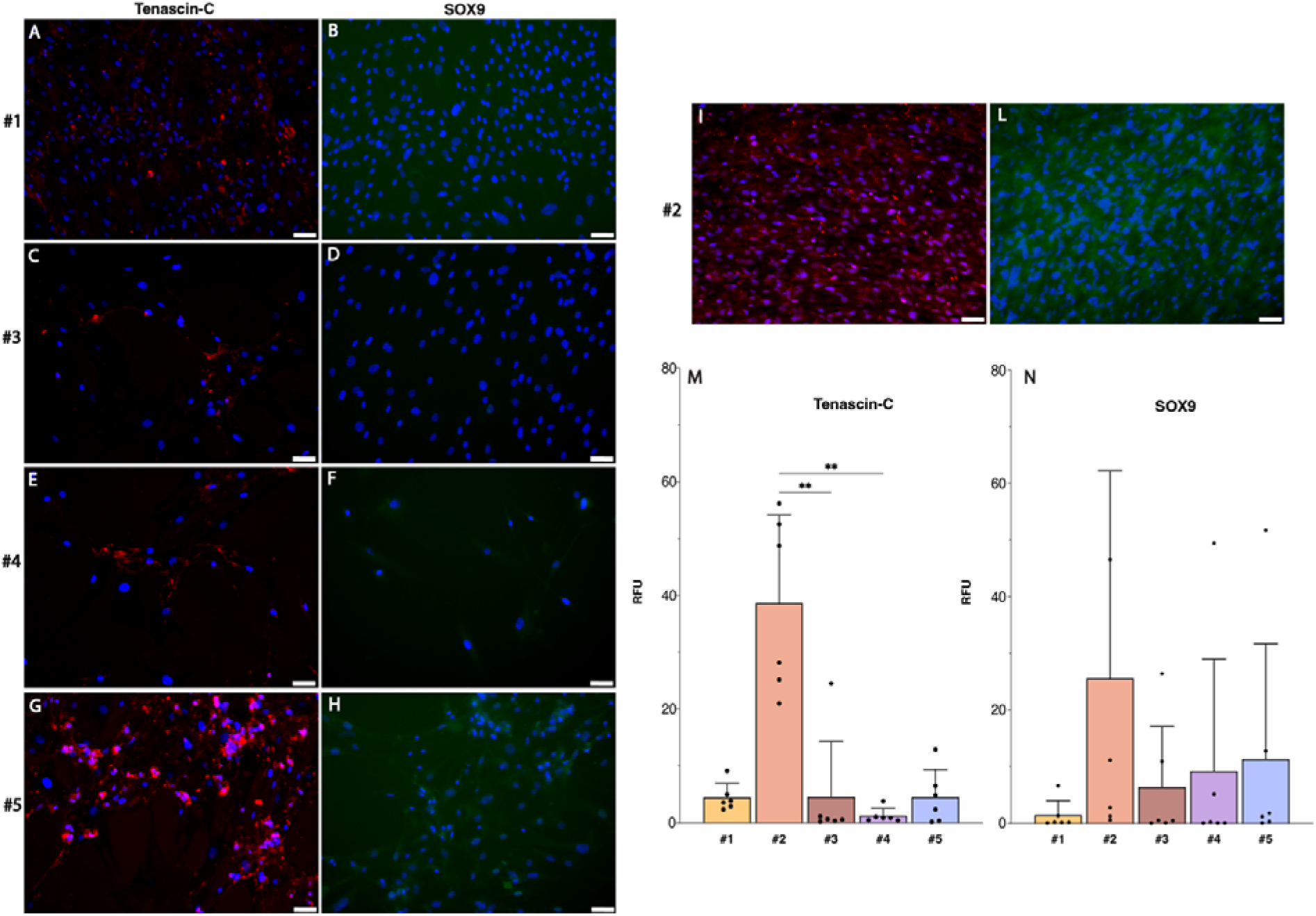
Immunofluorescence analysis of TNC (left column) and SOX9 (right column) expression in hTCs, showing the time-dependent effects of growth medium formulation on the proliferation rate, and tenogenic differentiation of hTCs in 2D culture at day 14. Scale bar = 50 μm. Quantification of TNC (M) and SOX9 (N) expression at day 14. Quantitative analysis was performed via the non-parametric Kruskal-Wallis test. For the quantification, six images were randomly acquired over three independent replicates, (n = 6). Data are reported as mean ± S.D, ** = p < 0.01.

Moreover, the #2 medium was found to be the most performing among all media screened, based on a scoring system we accurately developed, assigning scores from 1 to 5 based on cell shape, cell density within the well, and matrix formation capacity (figure SI2).

### Rolipram delivery supports tenocytes phenotype and ECM production

A preliminary cellularization test of our scaffolds with SW-1353 cells imaged after nuclear staining with DAPI, showed a high-rate cell adhesion to the microfibrous architecture, as well as the inhibition of cell sprouting from the scaffold (figure SI 3). The results from quantitative fluorescence performed over our scaffolds confirm such a protective effect toward resident TCs, as they produced a significantly higher amount of type I collagen when cultured up to 14 days within Rolipram loaded scaffolds (PLA/Rol/Coll), compared to both PLA, and PLA/Col samples (figure 9).

**Figure 9.**
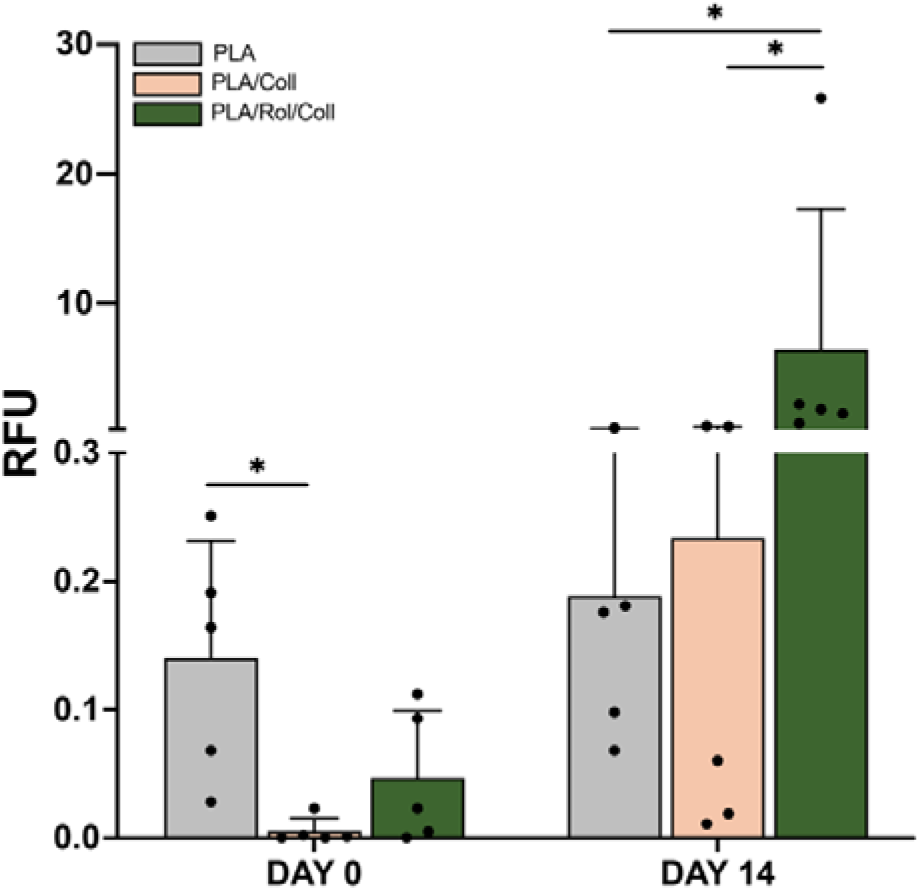
Type I collagen quantitative fluorescence performed via confocal microscopy on scaffolds cellularized with hTCs at day 0, and day 14. Mann-Whitney non-parametric test. For the quantification, five images were randomly acquired over three independent replicates, (n = 5), *p < 0.05.

The health of resident hTCs was further confirmed by the results from quantitative analyses for TNMD, which is among the stronger positive markers for the TCs functionality [29]. Remarkably, the TNMD showed a characteristic pericellular distribution to fill the gap between adjacent hTCs nuclei (figure 10).

**Figure 10.**
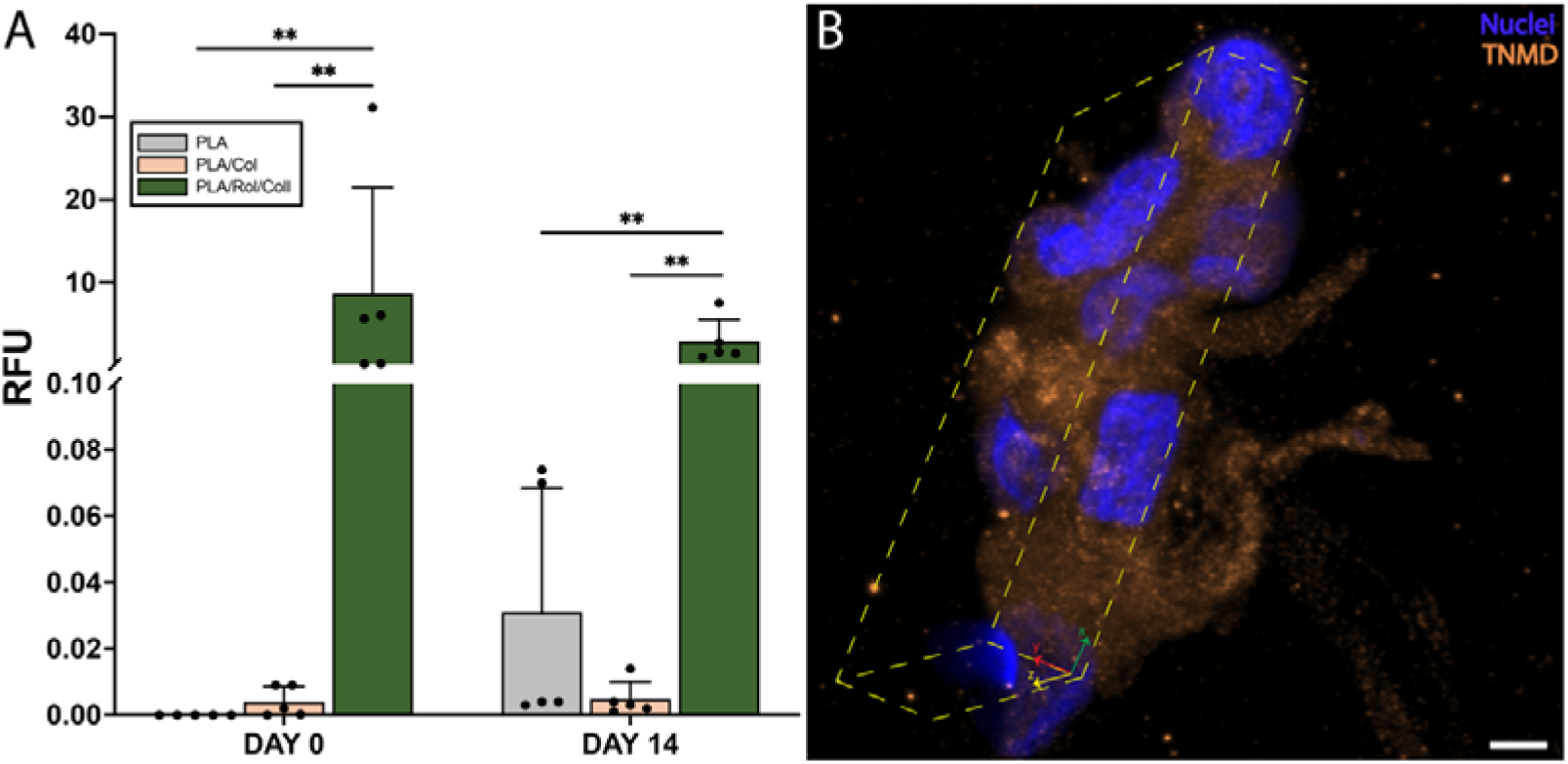
TNMD quantitative fluorescence performed via confocal microscopy on scaffolds cellularized with TCs at day 0, and day 14. Mann-Whitney non-parametric analysis. For the quantification, five images were randomly acquired over three independent replicates (n = 5), *p < 0.05, **p < 0.01 (A). Pericellular TNMD distribution in the new deposited ECM. Scale bar = 20 µm (B).

### Tendon-like electrospun scaffolds support cell growth and viability

Qualitative morphological analyses were conducted on scaffolds cellularized with hTCs and imaged on histological samples stained with H/E after 14 days of culturing (figure 11). Interestingly, a population of ECM-producing tenocytes is noticeable in each scaffold (figure 11, green arrows), as well as a cell distribution both in the outer and the inner portion of the scaffolds (figure 11, light blue arrows).

**Figure 11.**
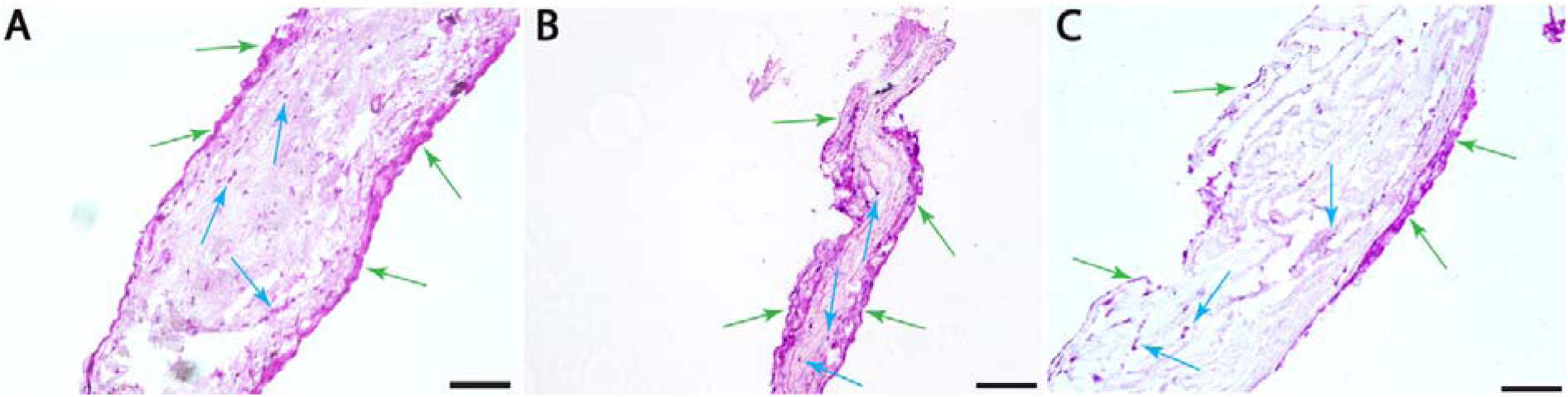
Hematoxylin/Eosin (H/E) staining of longitudinal sections of PLA (A), PLA/Coll (B), and PLA/Rol/Coll (C) electrospun scaffolds seeded with hTCs at day 14 of culture. Green arrows = new deposited extracellular matrix (ECM). Light blue arrows = resident hTCs in the inner portion of the scaffolds. Scale bar = 150 µm.

## DISCUSSION

Tendons are anatomical structures composed of dense, connective tissues organized in bundles of closely packed collagen I fibers which are responsible for the tendons’ mechanical strength and resilience. Collagen fibers are immersed within a proteoglycan/water matrix accounting for 60% -85% of tendon’s dry mass [30].

Among injuries interesting tendons, ruptures (TRs) represent the most significant clinical challenge due to the limited self-healing capacity of tendons, primarily caused by their low vascularity and cellularity. Although surgical repair is the gold standard approach for restoring tendon integrity, it often leads to nonfunctional fibrotic tissue that lacks the native mechanical resistance required for joint function. In this context, tissue engineering (TE) has emerged as a promising alternative, aiming to improve surgical outcomes or address early-stage tendon injuries (TIs). In fact, over the recent decades significant efforts have focused on creating hybrid scaffolds that integrate synthetic and natural polymers to achieve both mechanical stability and biocompatibility [31], [32].

In line with this trend, here we introduce an innovative and integrated strategy combining TE and drug delivery to prevent or mitigate the post-surgery tendon fibrosis (TF). We fabricated an engineered 3D electrospun (ES) scaffold with a tendon-like microarchitecture loaded with Rolipram and designed to locally deliver the drug with a prolonged release. Our overarching aim was to support both structural and functional tendon regeneration after a surgical repair.

In our study we used PLA to produce electrospun microfibers with an aligned pattern (figure 2A-D), and a high alignment degree (figure 2E), which closely mimics the natural arrangement and diameter of collagen fibers in native tendons [33]. Such a microarchitecture is capable of creating a favorable environment for TCs seeded onto microfibers, inducing a native-like cell elongation [32], [34].

The aligned pattern also contributes to the formation of micropores that allow passive diffusion of nutrients throughout the microfibers, ultimately supporting the overall cell viability and proliferation.

In addition, we implemented a collagen-coating on the microfibers’ outer surface to further enhance the overall cell viability in the long time [35], [36]. Surprisingly, the collagen coating also influenced the scaffolds’ microarchitecture inducing a self-assembling of small groups of microfibers into fascicle-like substructures. This effect is evident in images obtained from SEM analyses for collagen-coated samples (figure 2B, D), and compared with uncoated samples (figure 2A, C). We hypothesize that the collagen coating may randomly collect small groups of microfibers, promoting the self-assembly of those fibers, and stabilizing them into fascicular-like substructures. Remarkably, such substructures appear to be conserved also in 3D scaffolds that we fabricated to reproduce the native-like hierarchical architecture of tendons using a custom-designed bundle-making device (figure 1A). Such a device is designed to assemble microfibers into a helicoidal-like quaternary structure (figure 1B) in a repeatable and standardized manner. Such a fabrication technique allowed us to preserve the fascicle-like substructures, as is clearly visible in the digital rendering of the inner structure of our scaffolds analyzed by µCT (figure 3A-C).

We assessed the presence of collagen on the microfibers surface performing a qualitative FTIR-ATR analysis (IR), which confirmed the effectiveness of our collagen coating procedure as demonstrated by Amide I and Amide II collagen-specific peaks in IR spectra (figure 4). Precisely, the Amide I band related to C=O stretching vibrations in peptide bonds, displays a strong and well-defined peak at 1650 cm^-1^, a hallmark of type I collagen with its highly ordered triple helix structure [37], [38]. The Amide II band, linked to N–H bending and C–N stretching, further supports the identification of type I collagen by displaying a distinct peak at 1550 cm^-1^ reflecting the specific hydrogen bending within the triple helical structure.

Furthermore, the collagen coating also affected the overall mechanical behavior of our scaffolds. Under dry conditions, the incorporation of collagen led to a substantial increase in Young’s modulus, both for drug-free (from 102.93 to 140.09 MPa), and for drug-loaded (103.3 to 144.23 MPa) scaffolds. We attribute this behavior to a strengthening effect of collagen, whose adhesive properties restrict the fiber sliding during tensile testing. As a result, collagen-coated mats demonstrated reduced elongation at break and greater stiffness compared to uncoated samples, where fibers are free to slide on each other. These results are comparable to those reported by Lopresti and coworkers [39], who reported that chitosan-coated PLA ES mats demonstrated a significant modification of the mechanical properties. Conversely, the incorporation of Rolipram within our systems didn’t have any impact on the mechanical behavior, as demonstrated by the similar Young’s modulus of PLA and PLA/Rol samples, which was quantified in 102.93, and 103.3 MPa respectively. The collagen coating also influenced the scaffolds’ behavior when exposed to *in vivo*-mimicking wet conditions. In this case, both PLA and PLA/Rol mats exhibited a marked decrease in Young’s modulus and tensile strength, with about a 50% reduction compared to Young’s modulus registered in dry conditions for the same samples. Conversely, collagen-functionalized scaffolds maintained similar mechanical properties to those exhibited in the dry state, with only a slight decrease of Young’s modulus during tensile tests in wet conditions. We attribute such stiffness reduction to collagen hydration that induces a decrease of the collagen-fibers interactions, ultimately reducing the collagen adhesive effect. Furthermore, we believe that the higher temperature during wet testing (37°C) compared to dry testing (room temperature) promotes the mobility of the PLA backbone, exerting a synergistic effect on such a stiffness reduction. Overall, the mechanical properties depicted so far have a great analogy with those registered for the human tibialis posterior tendon [40], suggesting that our scaffolds are potentially compatible with a high-demanding environment as joints *in vivo*.

We next investigated how the collagen coating affects the drug release behavior. PLA/Rol scaffolds released significantly more drug compared to PLA/Rol/Coll when the values were normalized to the scaffold weight (figure 7). This observation is attributed to the presence of approximately 28 wt% collagen in the collagen-coated scaffold. Since Rolipram is only embedded within the PLA matrix, the overall amount of drug per unit weight of scaffold is consequently lower in the coated system, which explains the observed results.

To account for this difference, we also plotted the release profiles as M_t_/M∞, where M∞ represents the theoretical maximum amount of drug that can be potentially released from the system, assumed to be equal to the amount of Rolipram initially loaded into the PLA matrix. Even under this normalization, the release kinetics reveal that uncoated systems exhibit a faster release compared to their collagen-coated counterparts. Specifically, PLA/Rol scaffolds released approximately 60% of the loaded Rolipram within the first 4 hours, whereas collagen-coated scaffolds required 48 hours to reach the same percentage.

To quantitatively investigate the release kinetics and mechanism, the parameters *k* and *n* were determined according to the Peppas model [21]. The kinetic constant *k* was found to be 0.54 ± 0.035 for the PLA/Rol scaffold and 0.40 ± 0.06 for PLA/Rol/Coll, with a corresponding correlation coefficient of 0.957 ± 0.005 and 0.974 ± 0.011, respectively. The diffusion exponent *n* was 0.38 ± 0.036 for PLA/Rol and 0.40 ± 0.025 for PLA/Rol/Coll. These values indicate that Rolipram release from both scaffolds follows Fickian diffusion, as *n* is lower than 0.5 in both cases.

Although both systems exhibited the same release mechanism, the higher *k* value observed for uncoated scaffolds suggests a significantly faster release rate. This difference can be attributed to variations in structural and geometric features between the two systems. Specifically, *k* is known to be influenced by multiple factors, including matrix porosity, surface area, and characteristic diffusion path length [41], [42]. The presence of a collagen coating likely increases the fiber diameter and reduces surface porosity, which in turn extends the diffusion path and reduces the effective diffusion rate, as reported in similar systems [43], [44].

In addition to these encouraging findings, we moved forward by characterizing the biological behavior of our tendon-like scaffolds. One of the most common issues linked to the *in vitro* culturing of TCs in monolayer is their dedifferentiation toward phenotypes producing non tendon-specific ECM [45]. Hence, we first performed preliminary 2D culture of hTCs to choose the optimal culturing condition hampering the hTCs phenotype drift. With this aim, we have formulated 5 culturing media (conditions from #1 to #5, table 2) composed of different supplement types/concentrations and we employed them to culture hTCs in monolayer up to 14 days. Then, we assigned an overall score basing on three different morphological variables taking into account cell density (D), cells shape (S), and efficiency toward ECM production (M), which were evaluated at day 1, day 7, and day 14 of culture (figure SI2). We finally integrated the information from these analyses with the results obtained from the immunocytochemical studies involving positive, and negative TCs markers (TNSC, and SOX9 respectively, figure 8) that we have performed on day 14 for each condition. Intriguingly, both results converged toward the classification of condition #2 (figure 8 I-N) as the optimal culturing formulation, capable of supporting the hTCs growth without any interference with their native phenotype.

We exploited the results from those preliminary analyses to proceed with the tendon-like scaffold cellularization. We have preliminarily used a chondrosarcoma cell line (SW-1353) to evaluate the overall effectiveness of our cellularization protocol, as well as the real capability of our scaffolds to host, and support, a cell population. After we had obtained a positive set of results (figure SI 3), we moved forward by cellularizing our scaffolds with hTCs using condition #2 to culture the cellularized constructs for up to 14 days.

Despite the developmental derivation of TCs remains foggy [46], TCs are classified as specialized fibroblasts [47] since their direct differentiation from tenoblasts (which are tendon-specific fibroblasts) is supported by well-established evidence [4]. Also, it is well-known that healthy TCs produce an ECM rich in type I collagen hierarchically ordered, which ensures high elasticity, and resistance to tensile stresses characteristic of healthy tendons [48]. Rolipram exerts his specific antifibrotic effect by inhibiting the PDE4 in fibroblasts, ultimately leading to an increase of cAMP and a reduced production of collagen subfamilies, including the tendon-specific type I [49]. Hence, one of the most challenging aspects of our work was protecting the TCs we used to cellularize our scaffolds from un unwanted self-inhibition of the type I collagen production. Overall, data either from the type I collagen and the TNMD quantification, suggest that our scaffolds protect TCs, preserving their native phenotype and supporting the cell viability and functionality.

Hence, we analyzed each scaffold using a histomorphological approach based on hematoxylin/eosin staining (H&E) after the paraffin-embedding of our cellularized scaffolds. After the analysis of the results, we were impressed to notice hTCs localized both in the inner (light blue arrows, figure 11 A-C) and in the outer portions of each scaffold (green arrows, figure 11 A-C). The eosinophilic purple staining around outer cell nuclei suggests that our scaffolds supported the hTCs viability and contributed to maintain their native ability to produce new ECM up to 14 days. Intriguingly, such a cellular behavior is particularly evident for hTCs cultured within collagen-coated scaffolds, where the eosinophilic staining appears more intense. This aspect is not surprising, as collagen is among the most active ECM components driving the cells’ adhesion through discoidin domain receptors (DDRs) and integrin α1β1 interactions [50] [51].

## CONCLUSIONS

In this proof-of-concept study, we demonstrated the fabrication of a drug-loaded scaffold designed for tendon regeneration. This bioactive scaffold would facilitate tendon regeneration *in situ* by attracting host cells to the scaffold for tissue development while minimizing post-surgical fibrosis. The fiber alignment effectively mimicked the natural structure of tendon collagen, and sustained drug release, especially in the PLA/Rol/Coll condition, which persisted for up to 14 days, highlighting the efficacy of the collagen coating as drug-diffusion barrier. Collagen incorporation notably enhanced the mechanical properties of the 2D mats in dry conditions, resulting in a marked increase in elastic modulus and stiffness. Additionally, cellularization was successful across all conditions, with an improved cellular distribution observed in collagen-coated scaffolds. These findings underscore the beneficial impact of collagen coating in improving both mechanical performance and biological response, suggesting that these scaffolds are promising candidates for tendon repair applications. Nevertheless, these results represent early-stage observations and further biological characterization is essential to comprehensively assess the system’s regenerative potential. Future research should incorporate quantitative gene expression analysis (qPCR) to examine key tenocyte markers, and expand the panel of immunohistochemical assays to evaluate the expression of tendon-specific proteins within this 3D environment.

Overall, this preliminary investigation underlines the high potential of these tendon-like scaffolds, providing a basis for future efforts aimed at optimizing their design and efficacy in tendon repair strategies.

### Aknowledgments

Simona Campora is funded by the PON project on Research and Innovation 2014–2020 (Azione IV.4 -Contratti di ricerca su tematiche dell’Innovazione—B75F21002190001). The authors acknowledge the “Biomaterials analysis and preparation” laboratory of ATeN Center for the electrospinning equipment.

