## Supplementary figures and images for "Engineered collagen-coated scaffolds for tendon regeneration: a multifunctional drug delivery approach"

### Supplementary figure 1

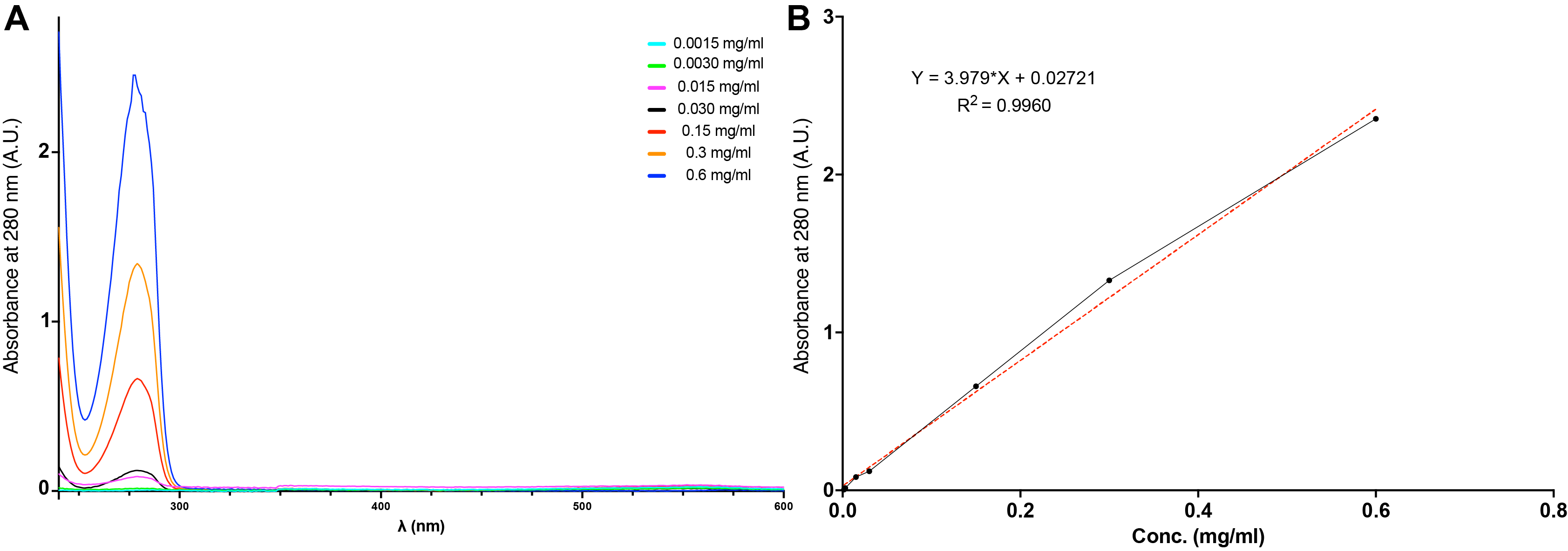

### Supplementary figure 2

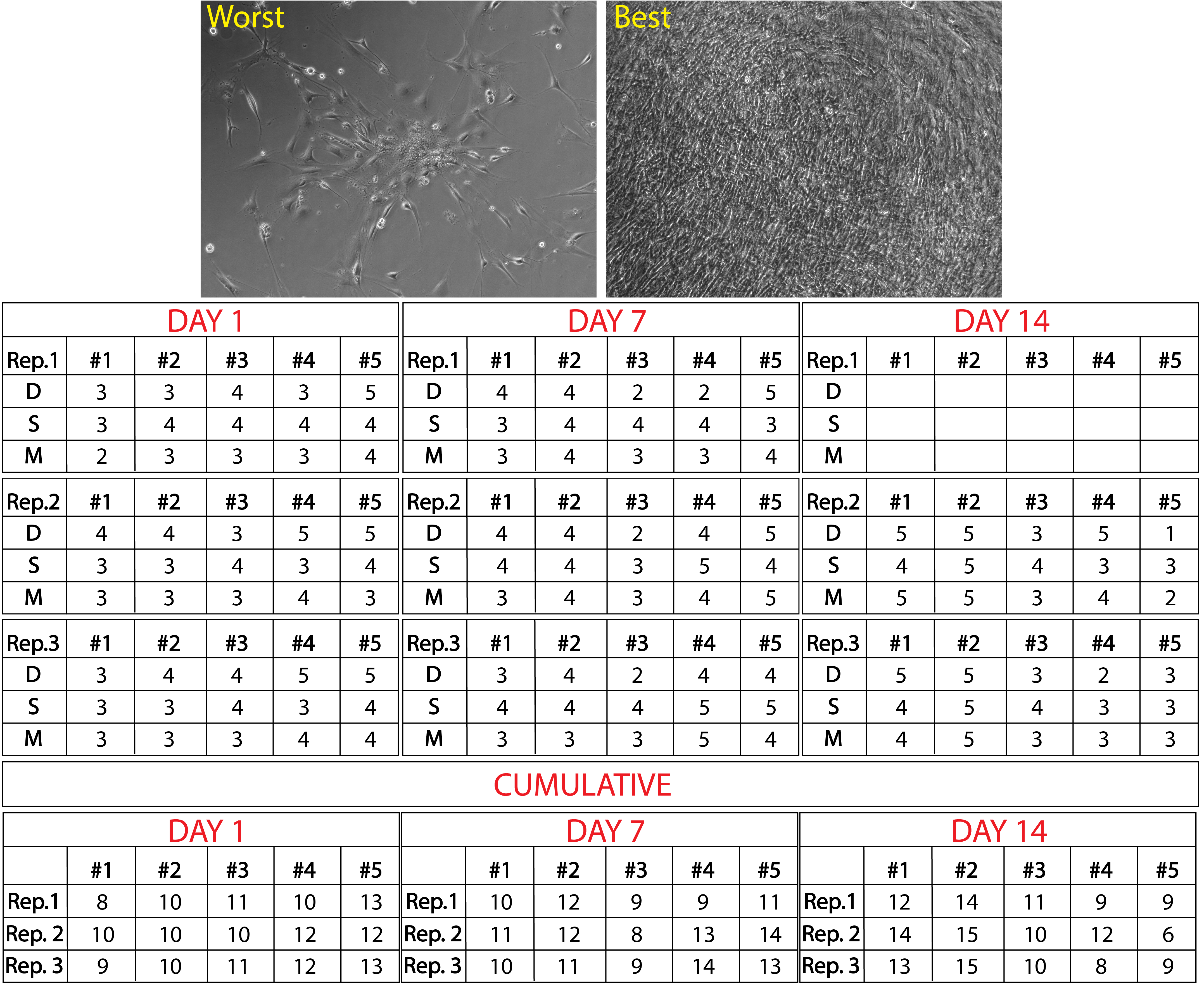

### Supplementary figure 3

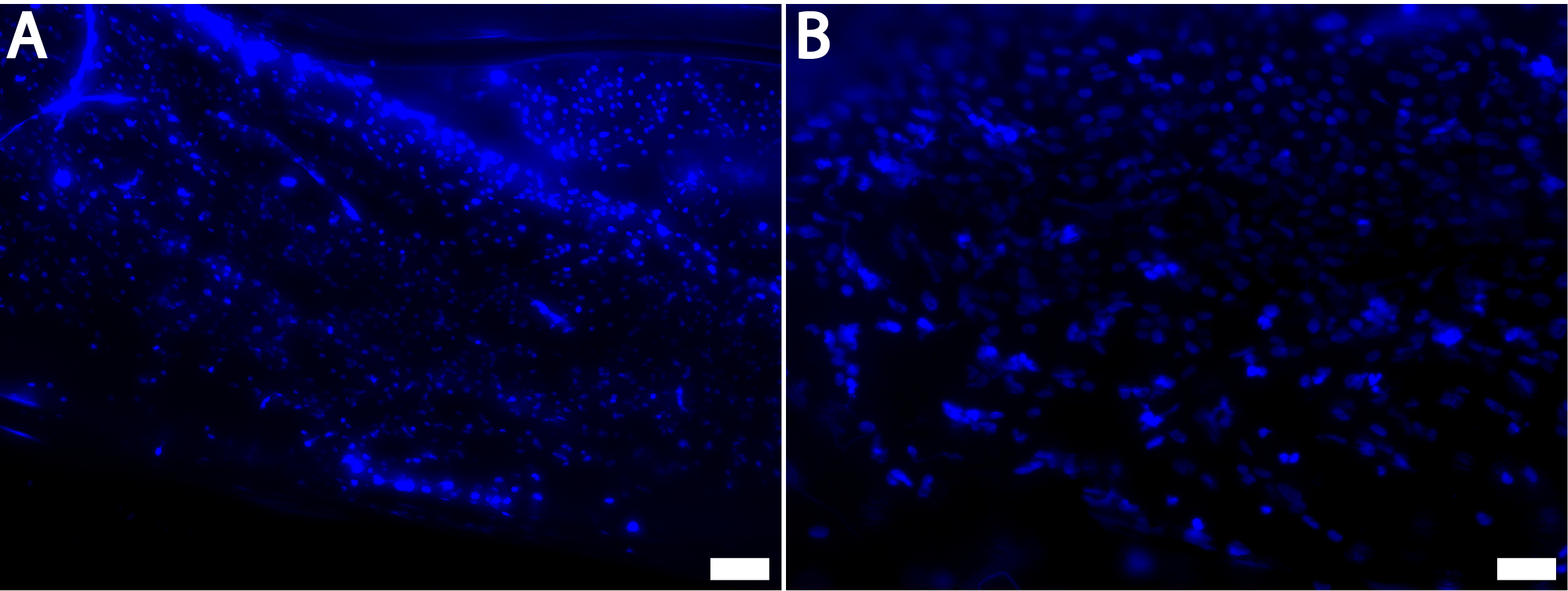
